# Automated anaerobic cultivation and phenotyping of diverse chain-elongating bacteria

**DOI:** 10.64898/2026.10.28.762647

**Authors:** Joel P. Howard, Christopher E. Lawson

**Affiliations:** Department of Chemical Engineering & Applied Chemistry, University of Toronto, Toronto, ON M5T 3E5, Canada

## Abstract

Anaerobic bacteria perform diverse metabolic functions with considerable biotechnological potential, yet their physiology remains poorly characterized because conventional anaerobic cultivation is low throughput. Here, we developed a Platform for Automated Cultivation of Microbial Anaerobes (PACMAN) and applied it to systematically characterize the physiology of diverse chain-elongating bacteria (CEB) that produce medium-chain carboxylic acids (MCCAs). High-throughput screening across nine CEB and 20 potential electron donors revealed extensive metabolic diversity, including previously unreported conversion of glycerol to octanoate by *Pseudoramibacter alactolyticus* and xylose to hexanoate by *Caproiciproducens* sp. 7D4C2. Comparison with genome-derived metabolic pathways further revealed widespread discrepancies between predicted and observed substrate utilization. Across diverse CEB and electron donors, decreasing the acetate-to-electron-donor ratio consistently shifted product distributions toward longer-chain carboxylates, revealing a general principle governing chain-length specificity, although maximum attainable chain length remained strain dependent. Finally, screening CEB across increasing carboxylate concentrations revealed substantial variation in MCCA tolerance, with Gram-positive MCCA producers exhibiting greater resilience than Gram-negative and butyrate-producing bacteria. Together, these findings reveal organizing principles underlying the metabolic versatility and product specificity of CEB and establish automated anaerobic phenotyping as a scalable approach for discovering microbial functions relevant to biomanufacturing.

## Introduction

Anaerobic microorganisms (anaerobes) are versatile biological catalysts that perform diverse metabolic functions^1^, including the conversion of complex waste feedstocks into high-value chemicals^2–4^. Anaerobic bioprocesses avoid the need for costly aeration and cooling^5^, while low biomass yields divert more carbon to products, making them economically attractive chemical production platforms^6^. Despite their importance, relatively few anaerobes have been isolated and far fewer have been functionally characterized^7–9^. As such, bioprocesses generally use model aerobic organisms^10^, which are often genetically engineered with considerable effort to perform reactions that uncharacterized anaerobes may already have evolved to carry out more efficiently.

An emerging anaerobic bioprocess is the conversion of organic waste into medium-chain carboxylic acids (MCCAs) using anaerobic chain-elongating bacteria (CEB)^2,11^. MCCAs are straight-chain monocarboxylates consisting of six (e.g. hexanoate – C6) to twelve (e.g. dodecanoate – C12) carbon atoms, and are used in fragrances, pharmaceuticals, feed additives, antimicrobials, lubricants, rubbers, dyes and liquid biofuels^2^. CEB synthesize MCCAs through the reverse β-oxidation (RBO) pathway, where acyl-CoA intermediates are elongated two carbons at a time by the addition of acetyl-CoA, before the cycle terminates via the transfer of the CoA moiety to a short-chain carboxylic acid (SCCA, one to five carbons), yielding a free carboxylic acid and a short acyl-CoA that can re-enter the cycle^2^. CEB have largely been studied as part of open microbial communities^2,12^, preventing detailed understanding of their metabolism as phenotypes cannot yet be reliably inferred from meta-omic analyses^13^. Accordingly, fundamental questions, such as what are the electron donors, product profiles, and MCCA product tolerances of different CEB are difficult to answer without pure culture experiments.

The first reported MCCA-producing CEB, *Clostridium kluyveri*, was described by Barker and Taha in 1942^14^, who found it could convert ethanol and acetate to hexanoate (C6). Between 1942 and 2010, only 10 additional MCCA-producing CEB pure cultures were discovered. Around 2010, the application of open microbial communities for MCCA production (termed, the carboxylate platform), spurred interest in MCCA-producing microbiomes^15,16^, resulting in an increased number of reported CEB in isolation with unique metabolisms. For example, *Megasphaera hexanoica* converts lactate to hexanoate and octanoate^17^, *Caproiciproducens galactitolivorans* converts galactitol to hexanoate^18^, and *Peptonealla octanoica* converts amino acids and lactate to hexanoate and octanoate^19^. Despite these studies, the pace of CEB physiological characterization relative to new isolates remains slow, limiting mechanistic advancement of chain elongation biotechnology.

Efforts to functionally characterize CEB, and anaerobes in general, are hindered by low-throughput cultivation techniques common in anaerobic microbiology. Traditionally, anaerobes are cultured in glass vials sealed with rubber septa (e.g. Balch or Hungate tubes)^20^, which ensures an anaerobic environment is maintained within the vial, but scales poorly to handling more than tens of cultures at a time. Microplate cultivations within an anaerobic chamber reduce working volumes to the microliter scale, allowing for hundreds to thousands of culture designs to be tested in parallel^20^, but manually assembling different combinations of strains, substrates and growth media is slow and prone to human error. These issues can be overcome using automation, including liquid handling robots, incubators, and plate readers that, when integrated, can allow for phenotypic measurements at temporal resolutions unattainable by a single human^20^. However, while automated cultivation under aerobic conditions is well established ^20,22,23^, few studies have developed automated platforms and workflows for high-throughput cultivation and phenotyping under anaerobic conditions, preventing dense characterization of poorly understood anaerobic microorganisms.

Here we develop a Platform for Automated Cultivation of Microbial Anaerobes (PACMAN) and use it to collect key phenotypic measurements across ten diverse CEB, including growth, substrate utilization, product formation, and MCCA product toxicity under varying electron donor and acceptor concentrations. Growth kinetics and end-point carboxylate product profiles across 1,245 strain-electron donor combinations revealed previously unreported CEB metabolisms, such as the conversion of glycerol into octanoate (C8) by *Pseudoramibacter alactolyticus* and the conversion of xylose into hexanoate (C6) by *Caproiciproducens* sp. 7D4C2. By varying the initial acetate concentrations of 571 CEB cultures, we found that a decreasing acetate-to-electron-donor ratio consistently shifted the product spectrum towards longer MCCA chain lengths. Finally, we challenged 1,457 CEB cultures with increasing concentrations of MCCAs and found that Gram-positive MCCA producers displayed greater MCCA resilience compared to their Gram-negative or SCCA-producing counterparts. Together, these results provide fundamental insights of the metabolic versatility of CEB, while offering scalable automated workflows for uncovering the phenotypic potential of anaerobes.

## Results

### A platform for high-throughput phenotypic characterization of anaerobic microorganisms

PACMAN consists of a liquid handling robot, a humidity-controlled incubation system, and a plate reader, all housed inside of an anaerobic chamber (Figure 1). All three devices are integrated and controlled using a custom Hamilton Venus method, enabling automated microplate transport within, and between, devices. Depending on the growth conditions, some anaerobes require over five days to reach saturation, posing potential evaporative challenges. When incubated in 96-well plates at ambient humidity, the liquid in unsealed plates completely evaporated less than three days (Supplemental Figure 2A). While gas-impermeable seals prevented evaporation, we found that condensation rapidly collected on the underside of seals, obscuring optical density measurements (Supplemental Figure 2B-D). Applying a 1°C temperature gradient across sealed plates prevented the accumulation of condensation and allowed for smooth optical density measurements (Supplemental Figure 2E). To our knowledge, temperature gradient incubation is not available in commercial multi-plate incubators, limiting throughput to a single plate. Instead, we found that incubating 96-well plates, covered with either breathable seals or lids, in a humidity-controlled incubator allowed for smooth optical density measurements and minimal evaporation.

**Figure 1.**
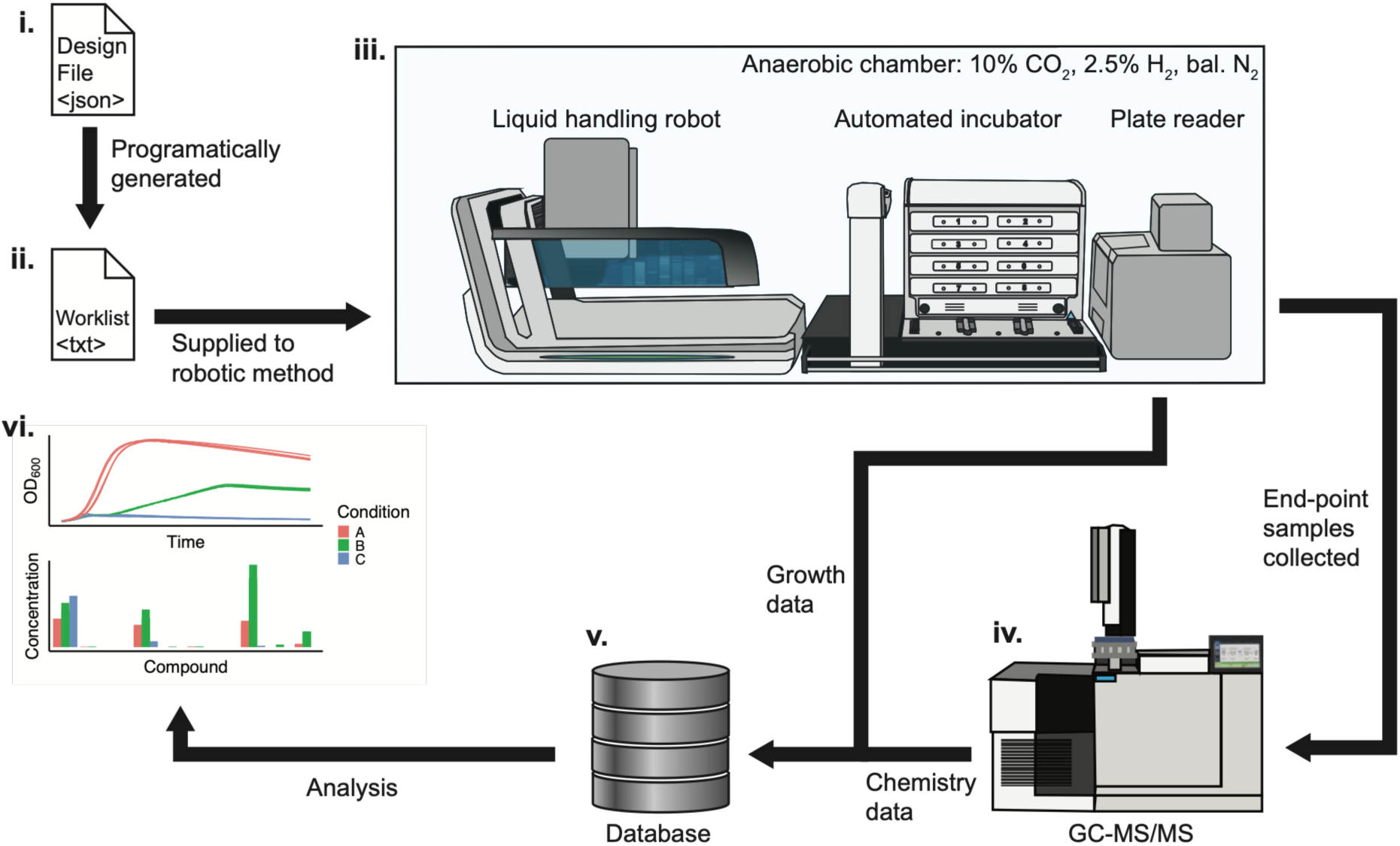
A high-throughput automated cultivation platform enables the functional characterization on anaerobic microorganisms. Schematic of the PACMAN. i) A json-formatted Design File, which specifies the strains, substrates, and media used for each design, as well as transfer volumes and liquid-property-specific transfer settings, is created by a user. ii) The Design File is parsed by a custom script which generates a tabular Worklist file. The Worklist is the set of liquid transfer instructions that the liquid handling robot uses to assemble the designed cultures in the Target Plates. iii) The automated devices in PACMAN consist of a liquid handling robot, an automated incubator and a plate reader housed inside an anaerobic chamber with a gas atmosphere of 10% CO_2_, 2.5 % H_2_, bal. N_2_. iv) Gas Chromatography–Tandem Mass Spectrometry is used to quantify S/MCCAs from end-point samples. v) Raw growth and chemistry data is parsed into tabular formats to create a phenotypic database. vi) Data is retrieved from the database for downstream analysis.

An experimental run using PACMAN begins with a user generated Design File, which specifies the combinations of media, supplements, and strains that are to be assembled (Figure 1). An upstream computational pipeline parses the information in the Design File to generate a Worklist, a list of transfer instructions for the liquid handler. Following the instructions described in the Worklist, the liquid handler transfers media, supplements, and strains from reagent reservoirs and 96-deep well plates into the on-deck 96-well Target Plate(s) (Supplemental Figure 1). To assemble samples in a reasonable amount of time, we implemented above-well multi-dispensing for substrates and strains. Allowing for the same pipette tip to dispense into multiple wells per aspiration reduces the time required to build plates by 2- to 4-fold and reduces the number of tips consumed by 3- to 21-fold, depending on the number of different strains, substrates, and media that are being combined (Supplemental Figure 3). We found that, when using multi-well dispense, as low as 10 μl of 200 g/l concentrated aqueous, volatile, or viscous stock solutions could be accurately dispensed into 96-well plates (Supplemental Figure 4).

Once a batch of Target Plates are built, the liquid handler automatically transfers them to the incubator and loads the next batch of Target Plates onto the deck from the Target Plate Stack. After all Target Plates are built and incubating, an integrated gripper continually cycles each Target Plate between its incubator slot and the plate reader, to track growth via optical density. Using this approach, PACMAN can cultivate 768 samples across eight Target Plates in a single experimental run. Once all samples have grown to saturation, supernatant is manually collected and C2-C8 carboxylate product profiles are measured (Methods).

### Profiling substrate-utilization and product-formation of chain-elongating bacteria reveals novel metabolic potential

With PACMAN established, we next identified 29 bacteria, spanning seven taxonomic families of the phylum Bacillota, reported to produce C6 (Figure 2A). To better understand the metabolic capabilities across diverse CEB, we selected nine of these CEB, across five taxonomic families, and a panel of 20 substrates that could potentially serve as electron donors for MCCA-production, including known electron donors for chain elongation, like ethanol, lactate, glucose, and fructose^16,24,25^, as well as less explored electron donors such as amino acids, xylose, and glycerol. We then tasked PACMAN with assembling the 180 strain-electron donor combinations and nine strain-basal media control combinations, totaling 1,245 cultures (N = 3-16 replicates per condition). Optical density at 600 nm (OD_600_) was tracked for all cultures, whereas GC-MS/MS throughput limited carboxylate analysis to a subset of 684 corresponding samples (N = 2-10 replicates per condition). We conducted all experiments in the same rich media (Methods) that supported growth of all CEB tested. Except for *R. intestinalis*, all CEB had some basal growth (OD_600_ > 0.1) on the media with no electron donor supplemented (Supplemental Figure 5), therefore we considered growth of a CEB on an electron donor as change in maximum OD_600_ greater than 0.25.

**Figure 2.**
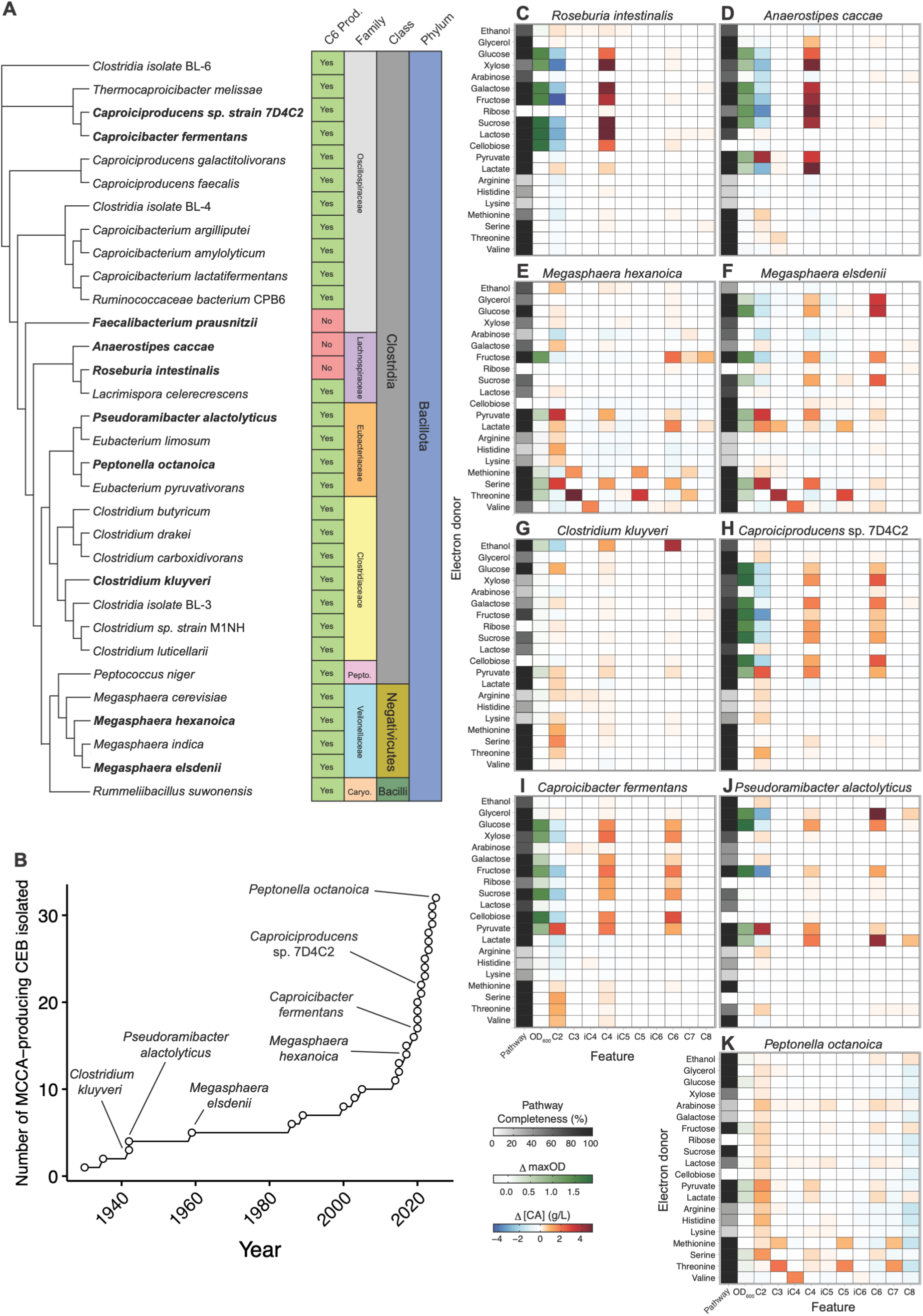
Substrate-utilization and product-formation profiles across diverse chain-elongating bacteria. (A) Hierarchical clustering of the 16S rRNA gene from all known hexanoate-producing CEB and selected butyrate-producing CEB. The names of CEB used in this study are bolded. Taxonomic family abbreviations: Pepto. – Peptococcaceae, Caryo. – Caryophanaceae. (B) The annual increase in the number of reported hexanoate-producing CEB, between 1930 and 2025. (C-K) Heatmaps showing the pathway completeness, change in maximum optical density and change in C2-C8 carboxylate product concentration in batch fermentations of nine CEB grown in the presence of 20 different electron donors. C2 – Acetate, C3 – Propionate, iC4 – Isobutyrate, C4 – Butyrate, iC5 – Isopentanoate, C5 – Pentanoate, iC6 – Isohexanoate, C6 – hexanoate, C7 – Heptanoate, C8 – Octanoate.

We found that, in the media and conditions used in this study (Methods), 75% of the electron donors could be utilized by at least one CEB for growth, with pyruvate being the most utilized, supporting growth of 89% of the CEB, whereas D-lactose, D-arabinose, L-arginine, L-histidine, and L-lysine were not utilized (Figure 2C-K). Under the conditions tested, *Anaerostipes caccae*, *Caproiciproducens* sp. 7D4C2, and *Megasphaera elsdenii* were the most versatile, utilizing 40% of the electron donors for growth, while *Clostridium kluyveri* was the most restricted, growing on only 10% of the electron donors. We were able to recapitulate well described electron donor-to-MCCA conversions, such as C6 production by *C. kluyveri* from ethanol^26^, C6 and C8 production by *P. alactolyticus* and *Megasphaera hexanoica* from lactate^27^, and C6 production by *M. elsdenii*, *M. hexanoica* and *Caproiciproducens* sp. 7D4C2 from fructose^25,28,29^. Notably, none of the cultures produced MCCAs with chain lengths greater than eight carbons, consistent with literature observations^27^.

Screening of strain-electron donor pairs also discovered unique substrates driving C6 and C8 production across several CEB, including glycerol, xylose, and amino acids. For example, both *P. alactolyticus* and *M. elsdenii* were found to utilize glycerol for growth and MCCA production, with *P. alactolyticus* producing high titers of C8. The SCCA-producer, *A. caccae*, also appeared to produce C4 using glycerol, but no growth above the basal level was observed. To our knowledge, this is the first time direct conversion of glycerol into MCCAs by CEB has been reported. To link these observations to specific genes, we reconstructed metabolic pathways from isolate genomes (Methods) to understand if phenotypes could be mapped to specific gene annotations. Despite only *P. alactolyticus* and *M. elsdenii* having clear growth and MCCA production on glycerol, we surprisingly found that seven of the nine CEB harboured all the necessary genes to oxidize glycerol into acetyl-CoA and produce carboxylates through RBO.

In addition to glycerol, *R. intestinalis* and *Caproicibacter fermentans* were observed to utilize xylose for growth and MCCA production, in agreement with previous studies^31–33^, as well as *A. caccae* and *Caproiciproducens* sp. 7D4C2, which has previously not been reported. Anaerobic xylose utilization begins with the isomerization of xylose to xylulose by xylose isomerase and phosphorylation of xylulose to xylulose 5-phosphate by xylulokinase. While our genome analysis did not identify a canonical xylose isomerase in any of CEB tested, it is possible that novel xylose isomerases in these genomes mediated xylose isomerization, as reported for other Bacillota^34^.

Several strains were also found to utilize amino acids as electron donors. *M. elsdenii*, *M. hexanoica,* and *Peptonella octanoica* displayed the most growth when cultured with no supplemental electron donor (Supplemental Figure 5), suggesting that they could utilize the amino acid rich extracts in the basal media. Indeed, they were the only CEB capable of utilizing at least one of the six amino acids tested. L-methionine and L-threonine can be oxidized to propanoyl-CoA, which can enter RBO instead of acetyl-CoA, resulting in the elongation of odd-chain carboxylates (e.g. propionate – C3, valerate – C5, and enanthate – C7)^19^. *M. elsdenii*, *M. hexanoica,* and *P. octanoica* harboured all the genes required to convert L-methionine and L-threonine into propanoyl-CoA, and we observed C3, C5, and C7 production when they were cultured in the presence of L-methionine or L-threonine*. M. elsdenii*, *M. hexanoica* and *P. octanoica,* were also found to utilize L-serine, in agreement with their genomic features. While *M. elsdenii* and *M. hexanoica* converted L-serine into C6, *P. octanoica* produced C4. *M. elsdenii* and *P. octanoica*, but not *M. hexanoica* which lacked a branched-chain amino acid aminotransferase, encoded the required genes to deaminate and oxidize L-valine into iso-butanoyl-CoA, which could enter RBO, or terminate in iso-butanoate (iC4). Interestingly, while none of these amino acid utilizing CEB showed increased growth when L-valine was present, they all produced iC4.

Overall, the utilization of an electron donor by CEB without a complete catabolic pathway was observed in 5% of the strain-electron donor combinations. This includes the observations of xylose and L-valine already mentioned, and galactose utilization by *Caproiciproducens* sp. 7D4C2, ribose utilization by *A. caccae* and *C. fermentans*, and sucrose utilization by *M. elsdenii*. CEB with complete catabolic pathways for electron donors that could not be utilized was more prevalent, being observed in 22% of the strain-electron donor combinations. Together, this highlights that database-informed metabolic pathway prediction does not always reveal the metabolic capacity of a microorganism.

### Acetate to electron donor ratios determine product chain length across diverse CEB and substrates

Controlling MCCA chain length allows for increased product specificity and decreases downstream separation costs in MCCA-producing bioprocesses. For a limited number of CEB, it has been shown that decreasing the initial acetate to electron donor ratio (A:Ed) shifts MCCA products towards longer chain lengths, increasing MCCA specificity^24,27^. To understand if this phenomenon holds across diverse CEB and electron donors, we used PACMAN to assemble 84 strain-A:Ed combinations, totaling 571 cultures (N = 3-28 replicates per condition). As before, we tracked the OD_600_ for all cultures and quantified end-point carboxylate concentrations for a corresponding subset of 317 cultures (N = 2-10 replicates per condition).

We found that across all MCCA-producer and electron donor combinations, decreasing A:Ed resulted in a consistent shift toward longer chain length MCCA products (Figure 3A-O). For *P. alactolyticus*, *M. hexanoica* and *P. octanoica*, while almost no C8 production was observed at high A:Eds (>1.5), as A:Ed decreased, the molar fraction of carboxylate products shifted towards larger proportions of C6 and C8. The highest C6 specificity (78 mol.%) was observed when *P. alactolyticus* was cultured in glycerol-containing media at an A:Ed of 0.47 mol acetate/mol glycerol, while for C8, the highest specificity (51 mol.%) was observed when *P. octanoica* was cultured in lactate-containing media at an A:Ed of zero mol acetate/mol lactate.

**Figure 3.**
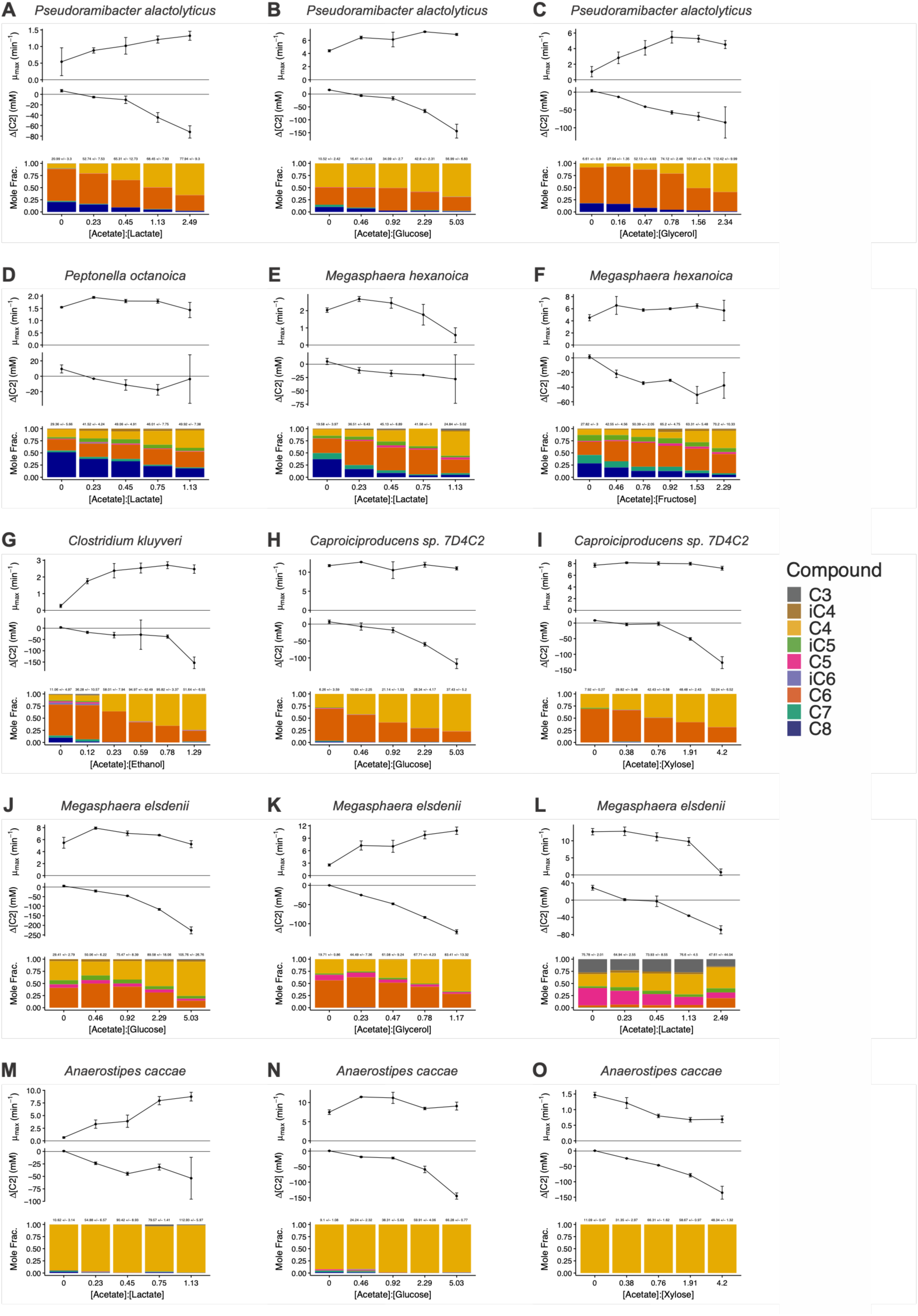
Electron donor and the acetate to electron donor ratio effects chain-length specificity and growth across CEB. Panels A-O show changes in growth rate (upper sub panel), acetate consumption (middle sub panel) and molar fraction of the C3-C8 carboxylate products (lower sub panel) for CEB across acetate to electron donor ratios. The mean total molar amounts of C3-C8 carboxylates +/−standard deviation are specified above each bar.

For the remaining MCCA producers, decreasing A:Ed resulted in increasing molar ratios of C6, but C8 production was never observed, even at A:Eds of zero. Similarly, for the SCCA-producer, *A. caccae*, none of the electron donor or A:Eds tested resulted in the production of carboxylates with chain lengths greater than four carbons, consistent with previous observations^27^. Therefore, although lowering the A:Ed shifts the product spectrum toward longer chain length MCCAs, certain strains are intrinsically limited in the maximum chain length that they can produce from RBO.

Changes in A:Ed appeared to have varying effects on strain growth rate. For *P. alactolyticus*, growth rate was positively correlated with increasing A:Ed of lactate and glycerol, while changes in A:Ed of glucose had little effect on the growth rate of *P. alactolyticus*. Increasing growth rate with an increasing A:Ed was also observed for *C. kluyveri* and ethanol, *M. elsdenii* and glycerol, and *A. caccae* and lactate. The growth rate of *Caproiciproducens* sp. 7D4C2 was constant across the A:Eds of glucose and xylose, but total carboxylate production decreased with A:Ed, suggesting that, at low A:Eds, *Caproiciproducens* sp. 7D4C2 disposed electrons on an alternative fermentation product. Lactate is a known fermentation product of *Caproiciproducens* sp. 7D4C2^25^. Lactate quantification of these same samples revealed that *Caproiciproducens* sp. 7D4C2, but not the other CEB, produced lactate in addition to S/MCCAs from sugar substrates (Supplemental Figure 6).

Changes in A:Ed did not affect the growth rate of *P. octanoica*, but any changes in growth rate may have been confounded by its high basal growth (Supplemental Figure 5). Similar near constant growth rates across A:Ed were observed for *M. elsdenii* and *M. hexanoica* growing on glucose and fructose, respectively. Interestingly, the growth rates of *M. hexanoica* and *M. elsdenii* decreased at high A:Eds of lactate.

### Gram-positive MCCA-producing CEB are more resilient to MCCA toxicity

MCCAs are antimicrobial compounds that inhibit bacterial growth by disrupting membrane integrity^35,36^, and are known to display differential toxicity to pathogens based on cell wall characteristics (e.g. Gram strain)^35^. Controlling MCCA concentration is essential in MCCA-producing bioprocesses to reduce the impact of product inhibition^12^. However, there are only scattered reports on MCCA toxicity for MCCA-producing mixed communities^16,37–39^, and fewer reports for individual MCCA-producing CEB^40^. To better understand the resilience of CEB to increasing concentrations of MCCAs, we selected six MCCA-producing CEB, two SCCA-producing CEB, and two Escherichia coli strains, and measured how their growth was affected by increasing initial concentrations of sodium hexanoate, sodium octanoate and sodium caprate (Figure 4). Using PACMAN, we cultivated 255 strain-electron donor-S/MCCA combinations across 1,457 cultures (2- to 18-replicates per condition).

**Figure 4.**
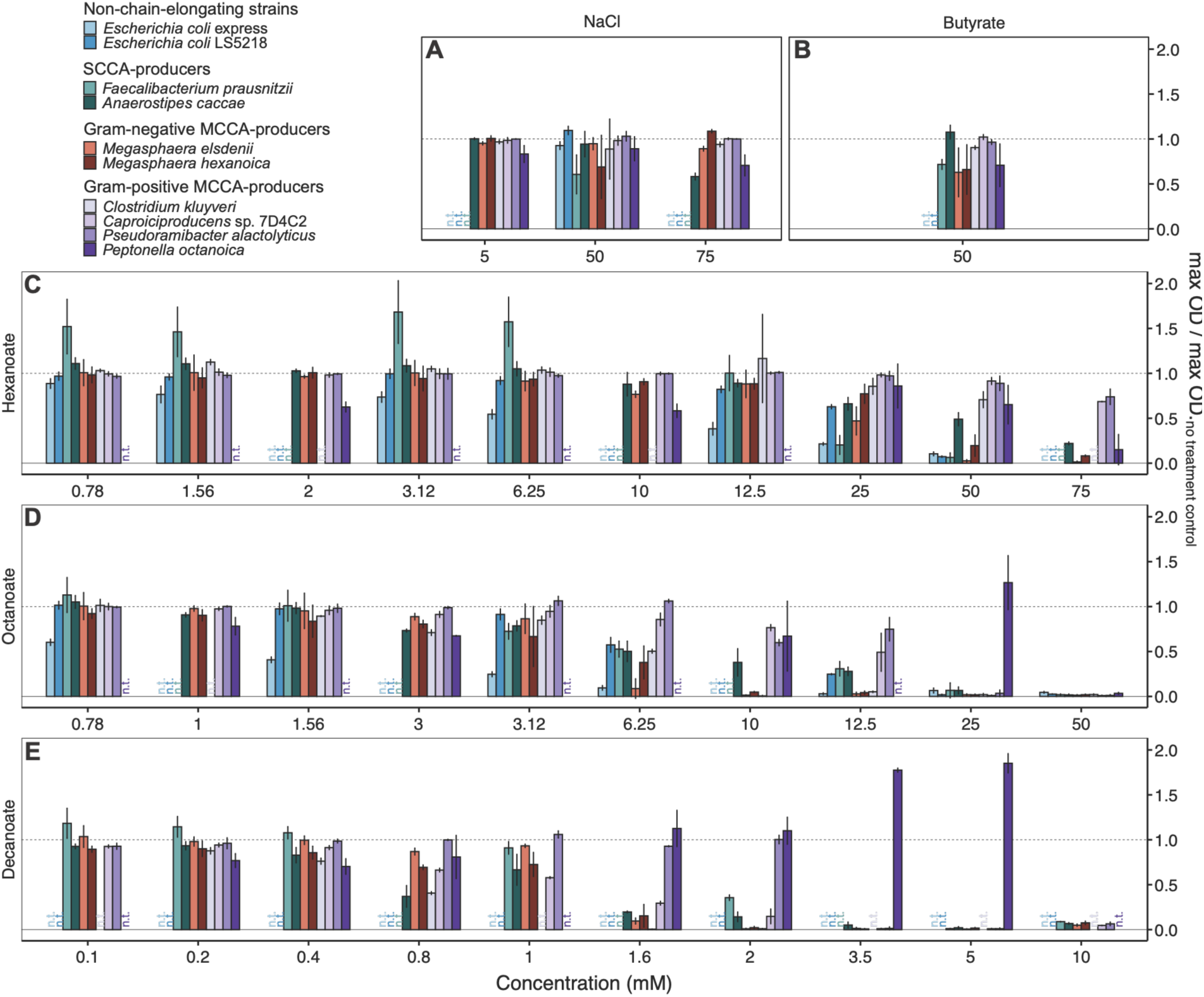
MCCA-producers display elevated resilience to product toxicity. The maximum optical density of eight CEB across increasing concentrations of (A) sodium chloride (NaCl), (B) butyrate, (C) hexanoate, (D) octanoate, and (E) caprate, relative to the maximum optical density of the un-supplemented control. Bars represent the mean value and error barse represent +/− the standard deviation.

As the MCCAs we exposed the cultures to were in the form of sodium salts, increasing MCCA concentration coincided with an increased sodium concentration. Therefore, we first wanted to determine if increasing sodium concentration would impact growth and confound MCCA toxicity measurements. We found that, for the majority of the strains tested, growth was unaffected after the addition of up to 50-75 mM of sodium chloride (Figure 4A). *F. prausnitzii* displayed reduced growth at sodium concentrations of 50 mM, while *P. octanoica* and *A. caccae* were sensitive to sodium concentrations of 75 mM in the absence of MCCAs.

When *Faecalibacterium prausnitzii, M. elsdenii, M. hexanoica*, and *P. octanoica* were subjected to an initial butyrate concentration of 50 mM, their growth decreased by 28%, 37%, 34%, and 29% respectively, while the other strains were unaffected (Figure 4B). *P. alactolyticus* and *Caproiciproducens* sp. 7D4C2, displayed the most resilience to hexanoate, respectively maintaining 74% and 68% of their maximum growth in the presence of 75 mM hexanoate (Figure 4C). *C. kluyveri* and *P. octanoica* were also tolerant to hexanoate, maintaining 71 and 65% growth, respectively, at a lower hexanoate concentration of 50 mM. While *M. elsdenii* and *M. hexanoica*, only maintained 3% and 2% growth at 50 mM, respectively, and did not grow at a hexanoate concentration of 75 mM. The SCCA-producer, *A. caccae* had a greater resilience to hexanoate than the *Megasphaera* strains, maintaining 49% growth at 50 mM and 22% at 75 mM. Despite appearing to increase growth at lower hexanoate concentrations, *F. prausnitzii* was the least resilient CEB to increasing concentrations of hexanoate, maintaining just 6% of its growth at 50 mM.

MCCA toxicity increased with increasing octanoate and caprate concentration, except for *P. octanoica*, where interestingly, an opposite effect was observed (Figure 4D,E). Indeed, the growth of *P. octanoica* was enhanced when treated with increasing concentrations of octanoate or caprate. The most resilient strains to octanoate and caprate were, *P. octanoica*, *P. alactolyticus*, *Caproiciproduces* sp. 7D4C2, and *C. kluyveri*. Grouping the MCCA-producers by Gram stain revealed that the Gram-positive MCCA producers were most resilient to increasing MCCA concentrations and chain lengths compared to their Gram-negative and SCCA-producing counterparts.

## Discussion

Automated cultivation has transformed the scale at which microbial phenotypes can be interrogated, but these approaches remain comparatively underdeveloped for anaerobic microorganisms. Here, we show that practical challenges associated with long-term anaerobic microplate cultivation can be overcome through humidity-controlled incubation, automated plate handling and repeated optical density measurements. Above-well multi-dispensing further reduced plate construction time by 2- to 4-fold and pipette-tip consumption by 3- to 21-fold, allowing nearly 800 anaerobic cultures to be assembled and monitored in a single experimental run. Throughput could be further increased by expanding incubator capacity beyond the eight plates used here. Unlike commercial phenotype screening approaches such as Biolog Phenotype MicroArrays, which rely on predefined assay panels and primarily measure metabolic activity through dye reduction, PACMAN permits user defined combinations of strains, media and supplements while directly measuring growth and enabling subsequent metabolite analysis. This flexibility is particularly important for fermentative anaerobes, where substrate utilization alone provides limited information about product formation.

Increasing cultivation throughput also revealed that chemical analysis, rather than cultivation, is now the major experimental bottleneck. GC-MS/MS analysis was conducted on a subset of samples, requiring approximately 46 days of GC-MS/MS runtime, while the analysis of all cultivated samples would have required an estimated 151 days. Coupling automated anaerobic cultivation with emerging high-throughput metabolite measurements, such as acoustic ejection mass spectrometry^41^, should therefore enable increasingly comprehensive interrogation of anaerobic microbial physiology.

High-throughput cultivation and phenotyping with PACMAN uncovered substantial variation in substrate utilization among CEB and identified metabolic capabilities not apparent from existing physiological descriptions or genome annotations. Several strains utilized diverse carbohydrates, organic acids, and amino acids, whereas others, such as *C. kluyveri*, remained highly specialized. Notably, *P. octanoica*, *M. hexanoica*, and *M. elsdenii* exhibited substantial growth in the basal medium and were the only strains to utilize the amino acids tested, consistent with our recent findings that these organisms directly couple Stickland fermentation with RBO^19^. This highlights the capacity of some CEB to exploit the complex mixtures of substrates likely encountered in organic wastes and gastrointestinal environments.

Glycerol represented a particularly important substrate expansion of known CEB metabolism. *P. alactolyticus* and *M. elsdenii* directly converted glycerol to MCCAs, with *P. alactolyticus* producing substantial C8. Glycerol is generated in large quantities as a biodiesel coproduct^42^, and is also released during triglyceride hydrolysis, making it relevant to lipid-rich organic wastes^43^. Previous mixed-culture studies have demonstrated MCCA production from glycerol, but have largely attributed this conversion to metabolic interactions in which glycerol fermentation generates intermediates such as ethanol, acetate, and butyrate that are subsequently used for chain elongation^44,45^. Our results instead demonstrate that individual CEB can directly couple glycerol metabolism to RBO. More broadly, the discovery of glycerol, xylose, and amino-acid utilization illustrates how the substrate repertoire of CEB extends beyond the small set of electron donors on which they have traditionally been studied. Importantly, predicted catabolic pathways frequently disagreed with observed phenotypes, reinforcing the value of systematic phenotyping for resolving this metabolic diversity.

Acetate availability emerged as a general determinant of product chain length across diverse CEB and electron donors. We recently showed that *P. alactolyticus* responds to acetate limitation during lactate fermentation by recycling acetate internally, enabling greater lactate utilization and production of longer-chain products at the cost of slower growth^27^. Here, we expand on this finding across glycerol, glucose, fructose, xylose, and ethanol metabolism, showing that decreasing acetate-to-electron-donor ratios consistently shifted product distributions toward longer-chain carboxylates across MCCA-producing CEB. Together, these findings suggest that acetate recycling is a broadly conserved strategy for balancing electron donor oxidation with chain elongation rather than a feature specific to lactate metabolism.

Surprisingly, the shift toward longer-chain products was not universally accompanied by slower growth, with growth penalties observed only for specific electron donors. This contrasts with *P. alactolyticus* growing on lactate, where the additional protein investment required for acetate recycling was proposed to constrain growth^27^, and suggests that protein allocation may not impose the same limitation across all substrates. Overall, these findings have important implications for process design; namely, acetate availability should be controlled relative to electron donor supply according to the desired product rather than simply maximized as a co-substrate. Strain selection remains equally important, as *P. alactolyticus*, *M. hexanoica*, and *P. octanoica* were the only strains observed to produce C8, whereas other CEB produced predominantly C6 even in the absence of added acetate.

MCCA toxicity represents another important constraint on chain elongation, and our results indicate that tolerance varies substantially among CEB. Carboxylic acids inhibit microbial growth through weak-acid uncoupling, while the greater hydrophobicity of MCCAs promotes partitioning into cell membranes, disrupting membrane integrity and fluidity^35,46,47^. Consistent with these mechanisms, MCCA inhibition is strongly dependent on pH and chain length and can constrain production in chain-elongation bioreactors.

Despite this toxicity, several MCCA producers remained remarkably resilient to C6–C10, with *P. octanoica*, *P. alactolyticus*, *Caproiciproducens* sp. 7D4C2, and *C. kluyveri* among the most tolerant organisms tested. This contrasts with the antimicrobial activity of MCCAs against other bacteria, including pathogens such as *Campylobacter jejuni*, which can be inhibited by C6–C10 at low-millimolar concentrations in a strongly pH-dependent manner^48^. The elevated tolerance of MCCA producers may reflect adaptation to their own metabolic products and provide an ecological advantage by allowing them to remain active at concentrations that inhibit competitors. Although the underlying mechanisms remain unknown, it is likely that differences in membrane composition, transport, or cellular stress responses are involved. Identifying these mechanisms could guide selection or engineering of strains with increased tolerance and ultimately enable higher MCCA titers.

## Methods

### Robotic platform

A Microlab Nimbus 4-channel liquid handling robot (Hamilton) was integrated with a BioSpa eight-position incubator (BioTek) and a Synergy H1 absorbance/fluorescence plate reader (BioTek) through Hamilton Venus (v4.6.0.8001) software. The three-component system was installed inside of an anaerobic chamber (COY, 2.5% H2, 10% CO_2_, bal. N_2_ atmosphere) with a HEPA-filter atmosphere purification system. The liquid handling robot, incubator and plate reader were controlled using the Hamilton Venus (v4.6.0.8001) and BioSpa OnDemand (v1.01.10) software.

### Bacterial strains and their preparation for storage

Eight MCCA-producing and two SCCA-producing CEB were selected. All strains, except *P. octanoica,* which was independently isolated by our group^19^, were obtained from DSMZ in the form of freeze-dried cultures. Strains obtained from DSMZ were revived by resuspension in the corresponding DSMZ-recommended medium under anaerobic conditions and statically incubated at 37 °C in anaerobic Hungate tubes. Actively growing cultures of all ten strains were mixed with anoxic glycerol to a final glycerol concentration of 15% and stored in O-ring sealed cryovials (Sarstedt, 72.694.006) at −80 °C.

### Preparation of bacterial strains for PACMAN

Precultures of each bacterial strain were prepared prior to each experimental run. To prepare the precultures, glycerol stocks were transported inside an anaerobic chamber (COY, 2.5% H_2_, 10% CO_2_, bal. N_2_ atmosphere) on a cold block (Corning, 07-210-050) that was stored at −80 °C. Frozen biomass from the glycerol stock was scraped into a Hungate tube containing sterile media using an inoculation loop. The revival media for each strain is specified in Supplemental Table 1. Frozen glycerol stocks were returned to −80 °C and inoculated tubes were statically incubated at 37 °C for a strain-specific duration specified in Supplemental Table 1. Prior to each experimental run, the preculture OD_600_ was normalized to 0.25 by manually diluting the actively growing cultures in mATCC2107 media (See below).

### Preparation of media and concentrated supplements used for PACMAN

All PACMAN experiments used modified ATCC Medium 2107 (mATCC2107), which was prepared 1.25X concentrated by dissolving 12.5 g tryptose, 12.5 g beef extract, 3.75 g yeast extract, and 6.25 g sodium chloride in 1000 ml of milliQ water, adjusting pH to ∼7.1 using sodium hydroxide, briefly bringing the media to a boil in the microwave before adding 0.31 g L-cysteine HCl, sparging the media with N_2_/CO_2_ gas, autoclaving in a sealed serum bottle (Chemglass, CLS-4217-05), and finally confirming pH to be ∼6.8. Concentrated stock solutions of supplements (e.g. electron donors, acetate, MCCAs) were prepared according to Supplemental Table 2. Briefly, the required mass or volume of the supplement was measured and dissolved in a minimal amount of milliQ water in a 50 mL conical tube. If required, the pH of the solution was adjusted to ∼6.8 using sodium hydroxide or hydrochloric acid and then diluted to the final supplement concentration using milliQ water. Tubes were added to the anaerobic chamber and allowed to passively degas for three days prior to starting a PACMAN experiment.

### A Platform for Automated Cultivation of Microbial Anaerobes (PACMAN)

For each experimental run, a tabular Design File, specifying the desired media-supplement strain combinations to be tested, was supplied to a custom computational pipeline that generated the Worklist, a tabular file of liquid transfer steps used by the liquid handler, and layouts for the supplement and strain stock 96-deep-well plates (Biobasic, BR581-96NS). A new worklist was generated before each run. Because, the worklist can be quite large (>1000 rows of instructions) and complex (specifies optimized pipette channel dispense order and multi-dispense sequences) manually writing each row of instructions is time consuming and would likely lead to human error. Therefore, custom scripts were used to parse the Design File and generate the worklist (Code available upon publication). Aliquots of each OD_600_ normalized strain were arrayed into the wells of a sterile 96-DWP (Culture Plate). Similarly, concentrated supplements were aliquoted into the wells of a second 96-DWP (Supplement Plate) and concentrated 1.25X mATCC2107 media was added to reagent reservoirs. The Culture Plate, Supplement Plate and reagent reservoirs were loaded onto the deck of the LHR. Up to three lidded 96 WPs (Greiner, 655161) (Target Plates) were initially loaded onto the deck, while the remaining lidded Sample Plates were loaded in a custom 3D-printed plate stack. A total of eight Target Plates, the capacity of the incubator, could be processed per experimental run. Finally, a rack of 300 ul CO-RE tips (Hamilton, 235985) was added to the deck of the LHR and the required number of additional racks, calculated by the Hamilton Venus method at runtime, was loaded onto the tip stack.

The liquid handler automated the construction of the cultures in the Target Plates according to the Worklist. First, channel grippers on the liquid handler automatically de-lidded the on-deck Target Plates. Next, 200 μl of 1.25X mATCC2107 media was transferred to each well of the de-lidded Target Plates. Then, Worklist-specified volumes of supplements were transferred from the Supplement Plate to specified wells of the Target Plates and each well was diluted with water to 240 μl. Finally, 10 μl of OD normalized bacterial cultures were transferred from the Culture Plate to specified wells of the Target Plates for a final culture volume of 250 μl. The liquid handler mixed the contents of Supplement Plate and Culture Plate wells prior to aspiration. To minimize the number of tips and time required to build a plate, supplements, strains, and water were dispensed into the Sample Plates from above the liquid level, allowing for multiple dispenses from the same pipette tip into various wells. To improve volumetric transfer accuracy, each channel aspirated 20 μl more than required and immediately dispensed 10 μl back into the well that it aspirated from, while the final 10 μl left in the tip was discarded when the tips were ejected into the tip waste. Once all cultures of the on-deck Target Plates were built, the plates were automatically re-lidded and transferred by the liquid handler to the incubator, which was set at 37 °C and maintained a relative humidity between 80-90%. Then the next batch of plates was automatically transferred from the Plate Stack onto the deck of the liquid handler and cultures were constructed as described above.

This was repeated until all Target Plates were built and incubating. Target Plates were automatically transported from the incubator to the plate reader, at one-hour intervals, where they would be incubated at 37 °C with a 1 °C gradient across the plate for 2 minutes to remove any condensation that formed on the underside of the lid during the transfer. The Target Plate would then be shaken at 365 cpm for 2 minutes in double orbital mode to homogenize the culture before the OD_600_ of each well was measured. Incubations were performed until all wells reached saturation, after which the Target Plate was manually removed from the anaerobic chamber and immediately centrifuged at 2,204 xg for 20 min. Supernatant from each Sample Plate well was then transferred to the corresponding well in a clean 96WP (Assay Plate). Assay Plates were sealed using an adhesive foil seal (BioRad, MSF1001B) and stored at −20 °C for downstream chemical analysis.

### Short and medium chain carboxylic acid quantification

Short and medium chain carboxylic acids (collectively, carboxylic acids) were quantified using gas-chromatography tandem mass-spectrometry (GC-MS/MS). The supernatant collected in the Assay Plates was filtered using 0.22 μm filter plates (Millipore, MSGVS2210) into clean collection plates (Greiner, 655161). Filtered samples were diluted 50X and acidified, using an aqueous 1 M solution of formic acid, to a final formic acid concentration of 100 mM. Samples were analyzed using an Agilent 8890 gas chromatograph equipped with an Agilent J&W DB-FATWAX Ultra Inert column (Agilent G3903-63008) coupled to an Agilent 7000D triple quadruple mass spectrometer. A PAL RSI 120 Series 2 autosampler injected 0.3 μl of each sample, which was introduced in split-injection mode (10:1 split ratio). Helium was used as the carrier gas at a flowrate of 1 ml/min. The GC inlet temperature was 250 °C and the oven temperature program began at 80°C for 1 min, then ramped by 20 °C /min to 200°C, then ramped by 5°C /min to 210 °C, then ramped by 20 °C /min to 250 °C where it was held for 5 min. The mass spectrometer was operated in dynamic multiple reaction monitoring (dMRM) mode. Target analytes were ionized and fragmented by electron ionization. The precursor and product ions of target analytes were selected using MassHunter Optimizer v10.0 (Agilent). Carboxylate concentrations in each sample were calculated from standard curves containing equimolar concentrations of C1-C7 (Supelco, CRM46975) and C8 (Sigma-Aldrich, C5038) carboxylic acids. As with the samples, standards were acidified using an aqueous 1 M solution of formic acid to a final added formic acid concentration of 100 mM. Analysis was performed using MassHunter Quantitative Analysis software v10.2 (Agilent).

For each batch of GC-MS/MS samples, the concentration of samples with a signal to noise ratio less than three was considered to be indistinguishable from background noise and set to zero. Depending on the health of the GC column carryover from the previous the injection was observed to a varying extent in blank samples that were ran approximately every six injections. The carryover concentration of a compound was found to be linearly correlated with the concentration of the compound measured in the previous injection (data not shown). For each batch of GC-MS/MS samples, a linear regression of the carryover versus the concentration in the previous sample was calculated for each compound of the blank samples and used to calculate a carryover concentration for each of the samples. The corrected concentration of each sample was calculated by subtracting the concentration of the sample by its carryover concentration. The corrections ranged from 0 to 18 mM.

### Genomic analysis for electron donor catabolism using gapseq

Genomes of the nine bacterial strains used in **Figure 2** were obtained from the NCBI RefSeq database. Metabolic pathways were predicted using gapseq^49^, which identifies enzymes by BLASTP searches of each genome against a reference database of metabolic protein sequences. Pathway predictions were generated with the *gapseq find -p all* command. Resulting MetaCyc degradation pathways for each electron donor were retained. Reactions leading to acetyl-CoA were retained. If the pathway did not reach acetyl-CoA additional reactions were appended to connect the electron donor degradation to acetyl-CoA. For each strain, every reaction was manually inspected and curated, including manual searches in genome annotation files for genes pertaining to missing reactions.

## Supplementary Material

**Supplemental Figure 1.**
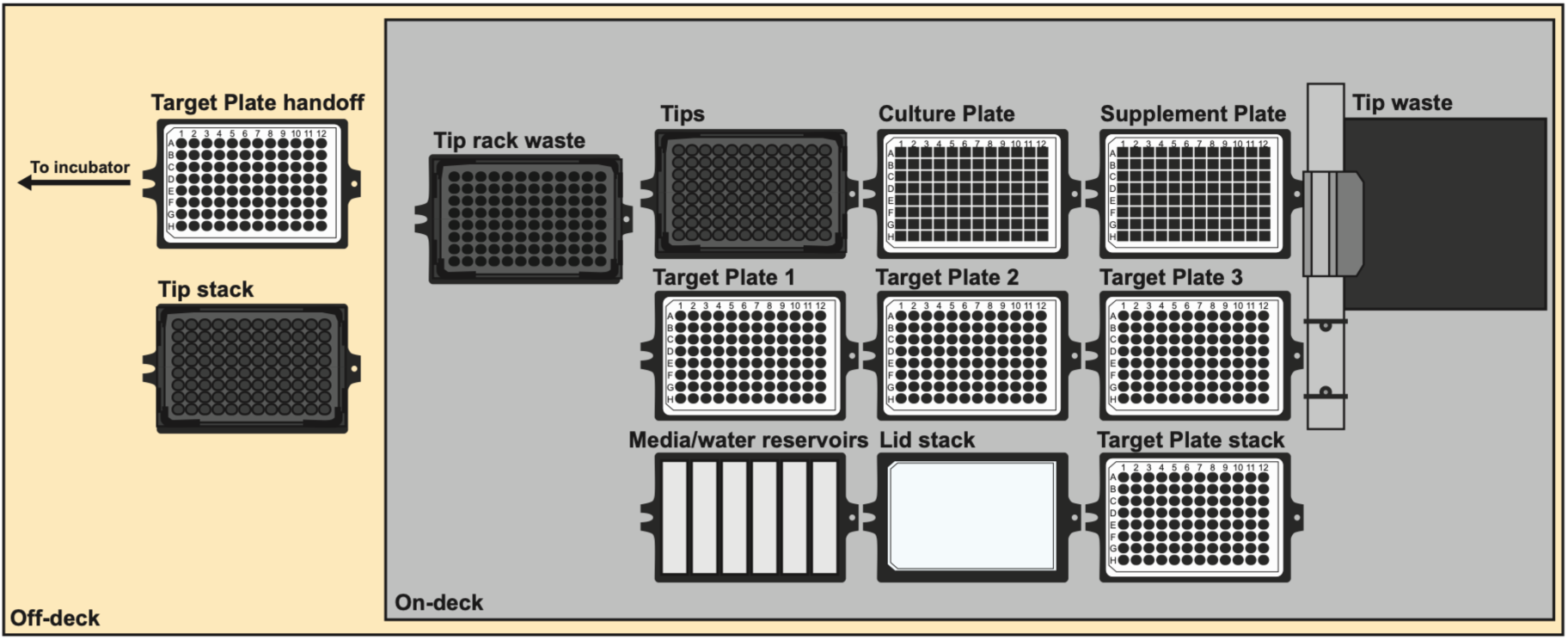
Deck layout of the PACMAN liquid handling robot.

**Supplemental Table 1.** Strains used in this study.

| Strain | Strain ID | Source | MCFA-producer | Revival Medium | Revival time (d) |
| --- | --- | --- | --- | --- | --- |
| *Anaerostipes caccae* | DSM 14662 | DSMZ | No | DSMZ<br>Medium 104 | 1 |
| *Caproicibacter fermentans* | DSM 107079 | DSMZ | Yes | DSMZ<br>Medium 1203a | 2 |
| *Caproiciproducens* sp. 7D4C2 | DSM 110548 | DSMZ | Yes | DSMZ<br>Medium 104c | 2 |
| *Clostridium kluyveri* | DSM 555 | DSMZ | Yes | DSMZ<br>Medium 52 | 3 |
| *Peptonella octanoica* | F2 | Food Waste<br>CSTR | Yes | DSMZ<br>Medium 104c | 4 |
| *Faecalibacterium prausnitzii* | DSM 107838 | DSMZ | No | DSMZ<br>Medium 1611 | 1 |
| *Megasphaera elsdenii* | DSM20460 | DSMZ | Yes | DSMZ<br>Medium 104 | 1 |
| *Megasphaera hexanoica* | DSM 106893 | DSMZ | Yes | DSMZ<br>Medium 1203a | 1 |
| *Pseudoramibacter alactolyticus* | DSM 3980 | DSMZ | Yes | DSMZ<br>Medium 455 | 2 |
| Roseburia intestinalis | DSM 14610 | DSMZ | No | DSMZ<br>Medium 1611 | 1 |
| *Escherichia coli* | LS5218 | Brian F.<br>Pfleger | No | LB | 1 |
| *Escherichia coli* | NEBExpress® | New<br>England<br>Biolabs | No | LB | 1 |

**Supplemental Table 2.** Supplements used.

| Compound | Form | Supplier | Product No | MW_Compound | MW_Form |
| --- | --- | --- | --- | --- | --- |
| cellobiose | D-(+)-CELLOBIOSE | Sigma | C7252 | 342.3 | 342.3 |
| lactate | SODIUM L-LACTATE, 98+% | Alfa Aesar | L14500 | 90.08 | 112.06 |
| pyruvate | SODIUM PYRUVATE | Aldrich | P76225 | 88.06 | 110.04 |
| lactose | D-LACTOSE MONOHYDRATE | Alfa Aesar | 36218 | 342.3 | 360.32 |
| sucrose | SUCROSE | Bioshop | SUC507 | 342.2965 | 342.3 |
| arabinose | L(+)-ARABINOSE | Bioshop | ARB222 | 150.13 | 150.13 |
| xylose | D(+)-XYLOSE | Sigma-Aldrich | X1500 | 150.13 | 150.13 |
| glucose | D-GLUCOSE ANHYDROUS REAGENT GRADE | Bioshop | GLU501 | 180.156 | 180.156 |
| galactose | D-(+)-GALACTOSE | Bioshop | GAL500 | 180.156 | 180.16 |
| fructose | D(-)-FRUCTOSE | Sigma | F2793 | 180.156 | 180.16 |
| glycerol | GLYCEROL | Bioshop | GLY002 | 92.09382 | 92.09 |
| ethanol | ETHANOL | Les Alcools d | P016EAAN | 46.068 | 46.068 |
| ribose | D(-)-RIBOSE | Sigma | R7500 | 150.13 | 150.13 |
| L-serine | L-SERINE | Bioshop | SER333 | 105.1 | 105.1 |
| L-threonine | L-THREONINE | Bioshop | THR002 | 119.12 | 119.12 |
| L-valine | L-VALINE | Bioshop | VAL201 | 117.151 | 117.151 |
| L-methionine | L-METHIONINE | Sigma | M5308 | 149.21 | 149.21 |
| histidine | L-HISTIDINE | Bioshop | HIS100 | 155.1546 | 155.16 |
| arginine | L-ARGININE | Bioshop | ARG006 | 174.2 | 210.7 |
| lysine | L-LYSINE MONOHYDROCHLORIDE | Bioshop | LYS101 | 146.19 | 182.65 |

**Supplemental Figure 2.**
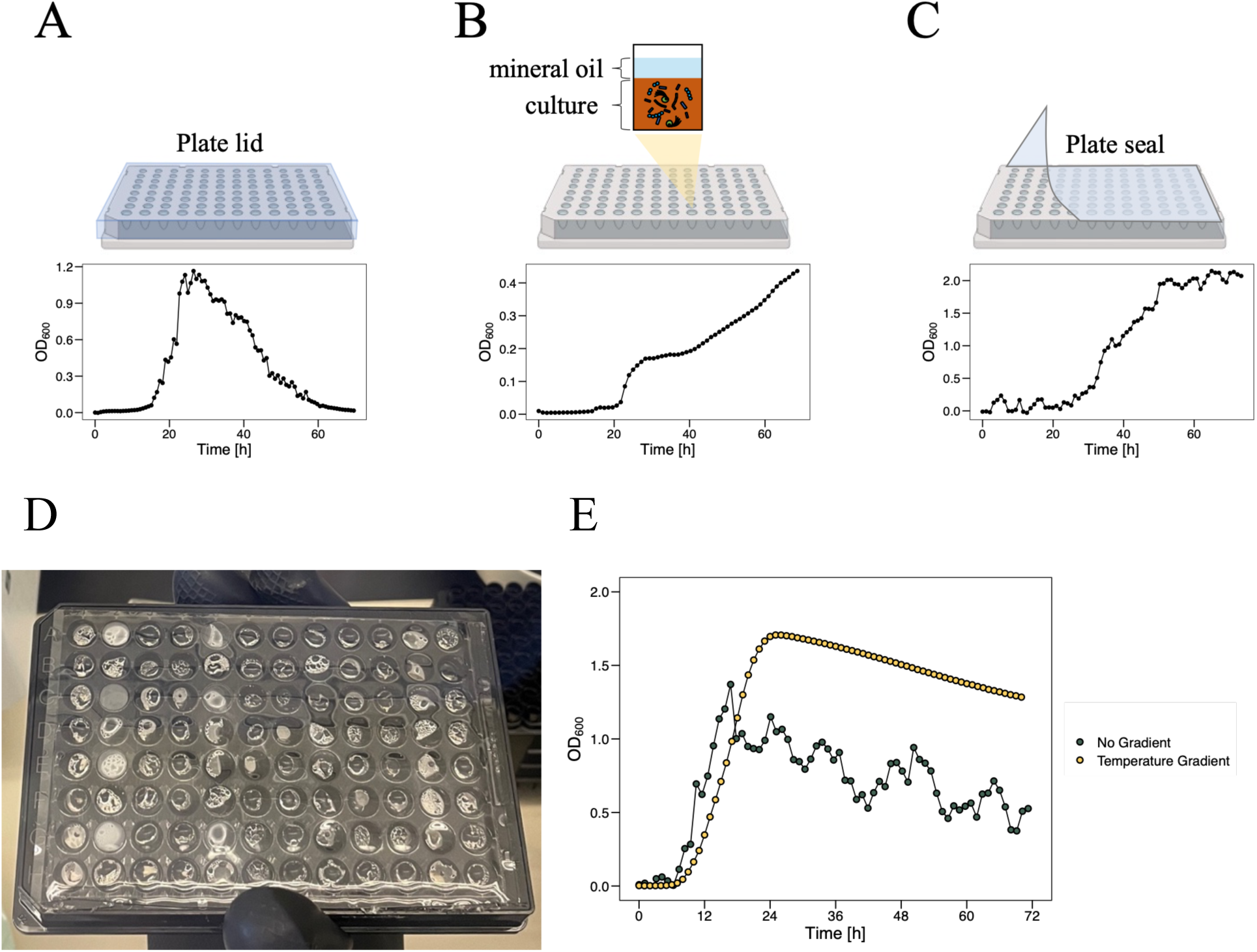
Evaluating different microplate covers to limit evaporation. A plastic microplate lid (A), a layer of mineral oil (B) and a PCR plate seal (C) were used to cover plates. Growth curves for *P. alactolyticus* growing in modified ATCC Medium 2107 are shown below the corresponding cover used to generate the data. (D) A microwell plate containing 200 μl aqueous culture in each well incubated at 37 °C without a temperature gradient. (E) Growth curves of *M. elsdenii* when incubated in a sealed microwell plate with (yellow) or without (green) a 1 °C temperature gradient between the bottom and top of the plate (top side hotter).

**Supplemental Figure 3.**
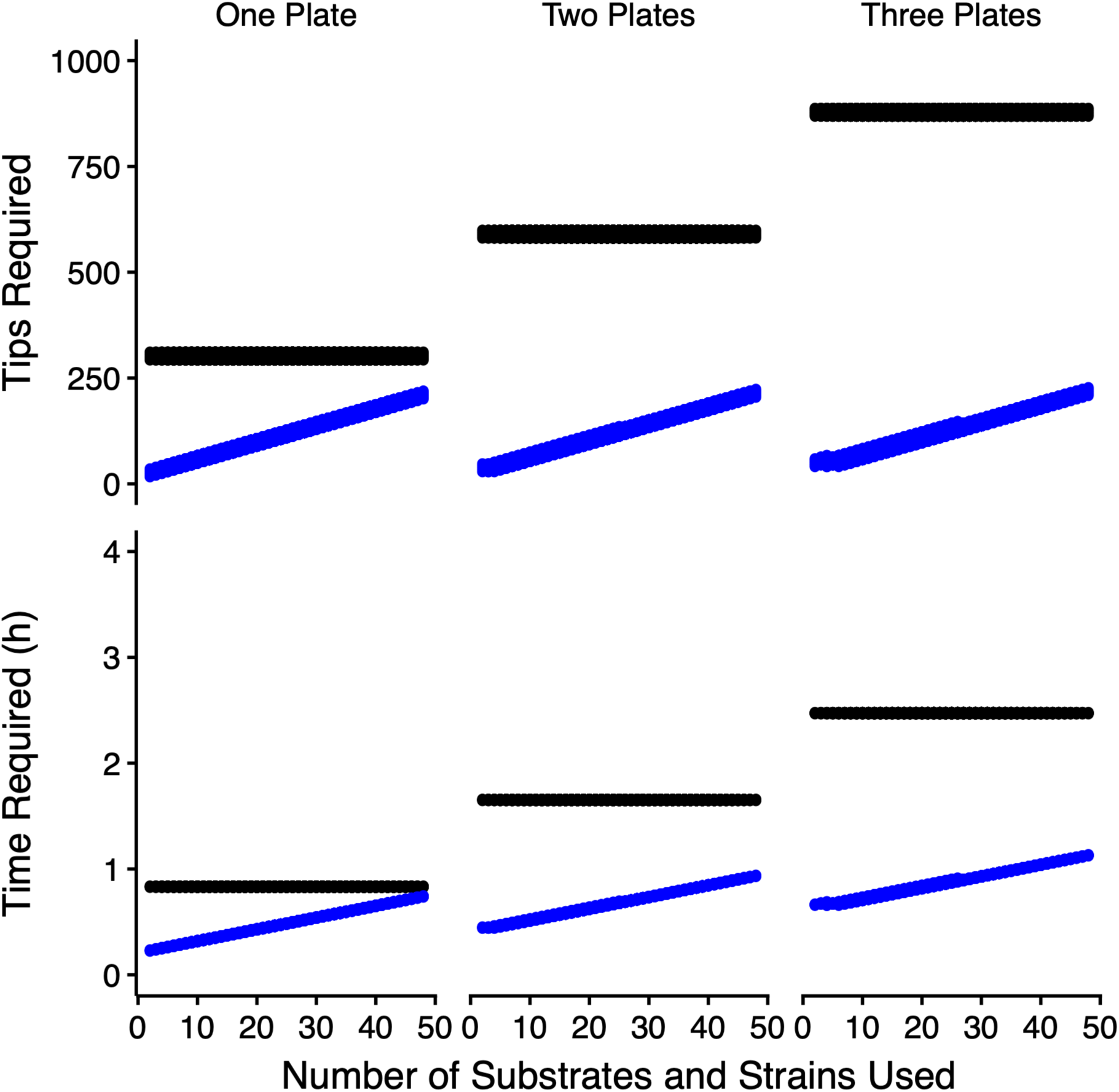
Multi-dispense provides significant consumable and time savings. Blue lines are multi-dispense and black lines are single-dispense. Columns separate data by number of Target Plates used in a batch (min one plate, max three plates).

**Supplemental Figure 4.**
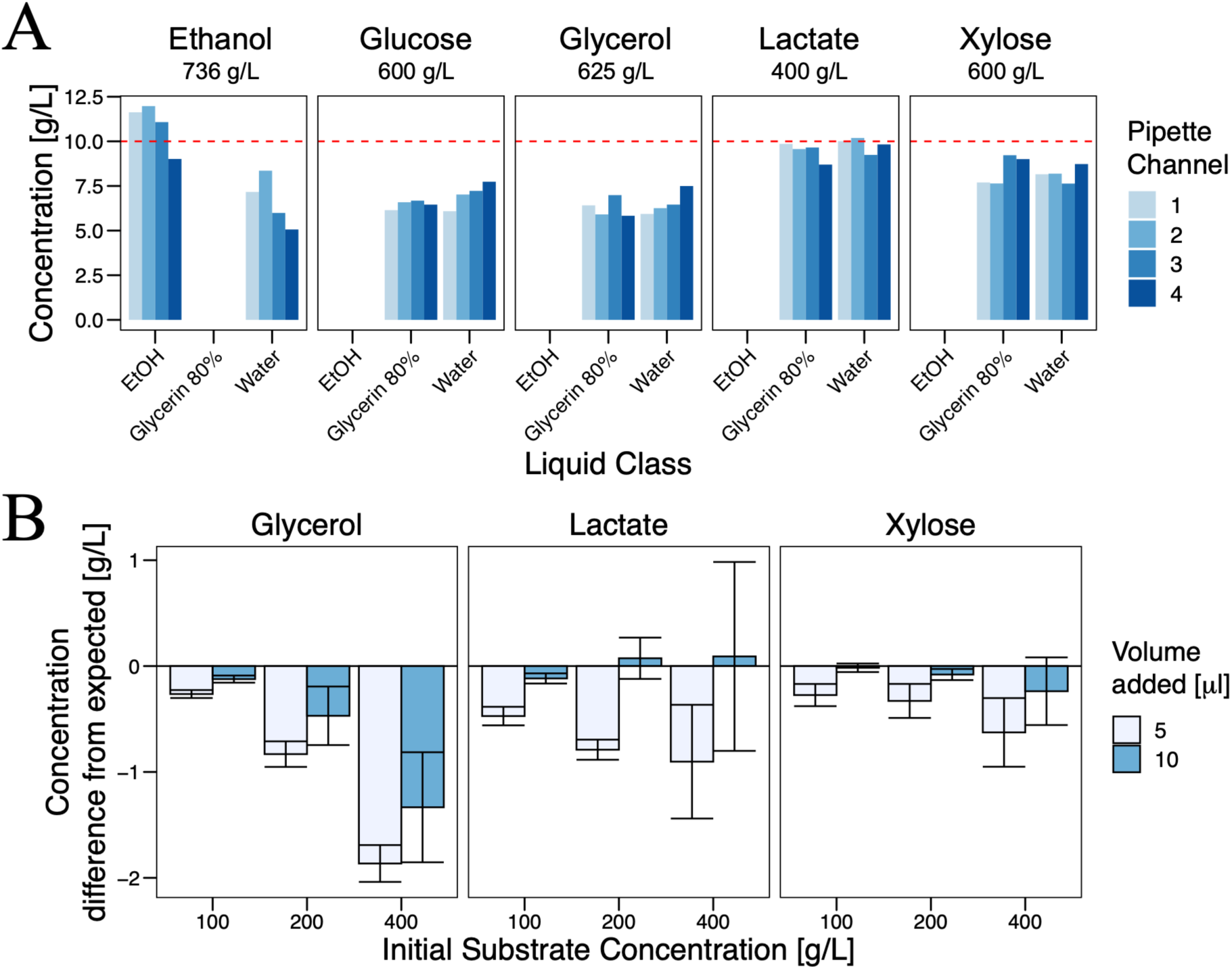
Evaluating the liquid transport accuracy of the LHR for volatile and viscous solutions. (A) The effect of different liquid classes – how the LHR transports the liquid – on transport accuracy for concentrated solutions. Dashed red line indicates expected concentration. Shades of bars indicate pipette channel used to make the transfer. N = 1 (B) Effect of initial substrate concentration and transfer volume on transfer accuracy. Bars represent mean difference from expected concentration and error bars are ± standard deviation. N = 4.

**Supplemental Figure 5.**
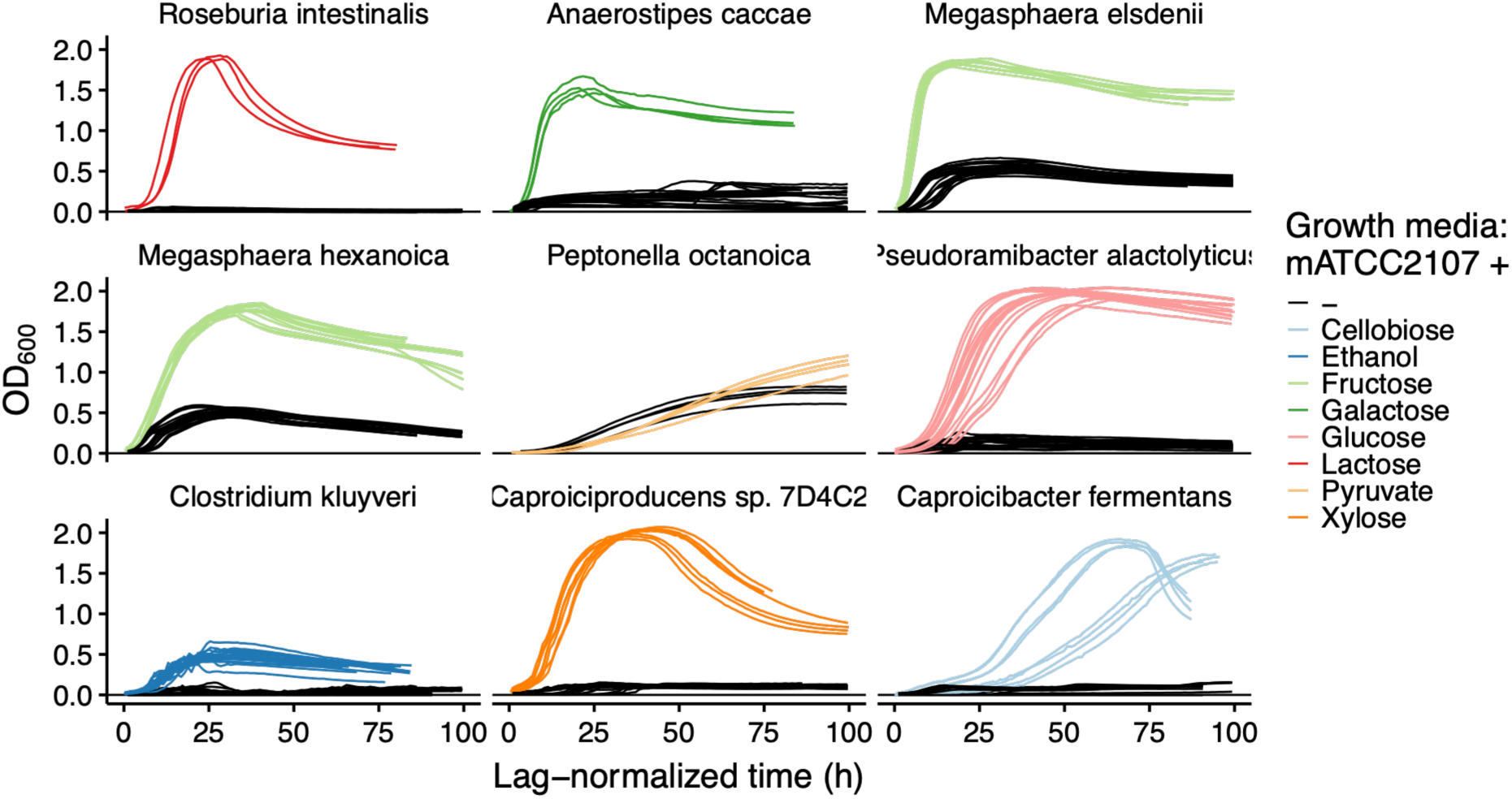
Growth of CEB on mATCC2107 with, or without, supplemented electron donor.

**Supplemental Figure 6.**
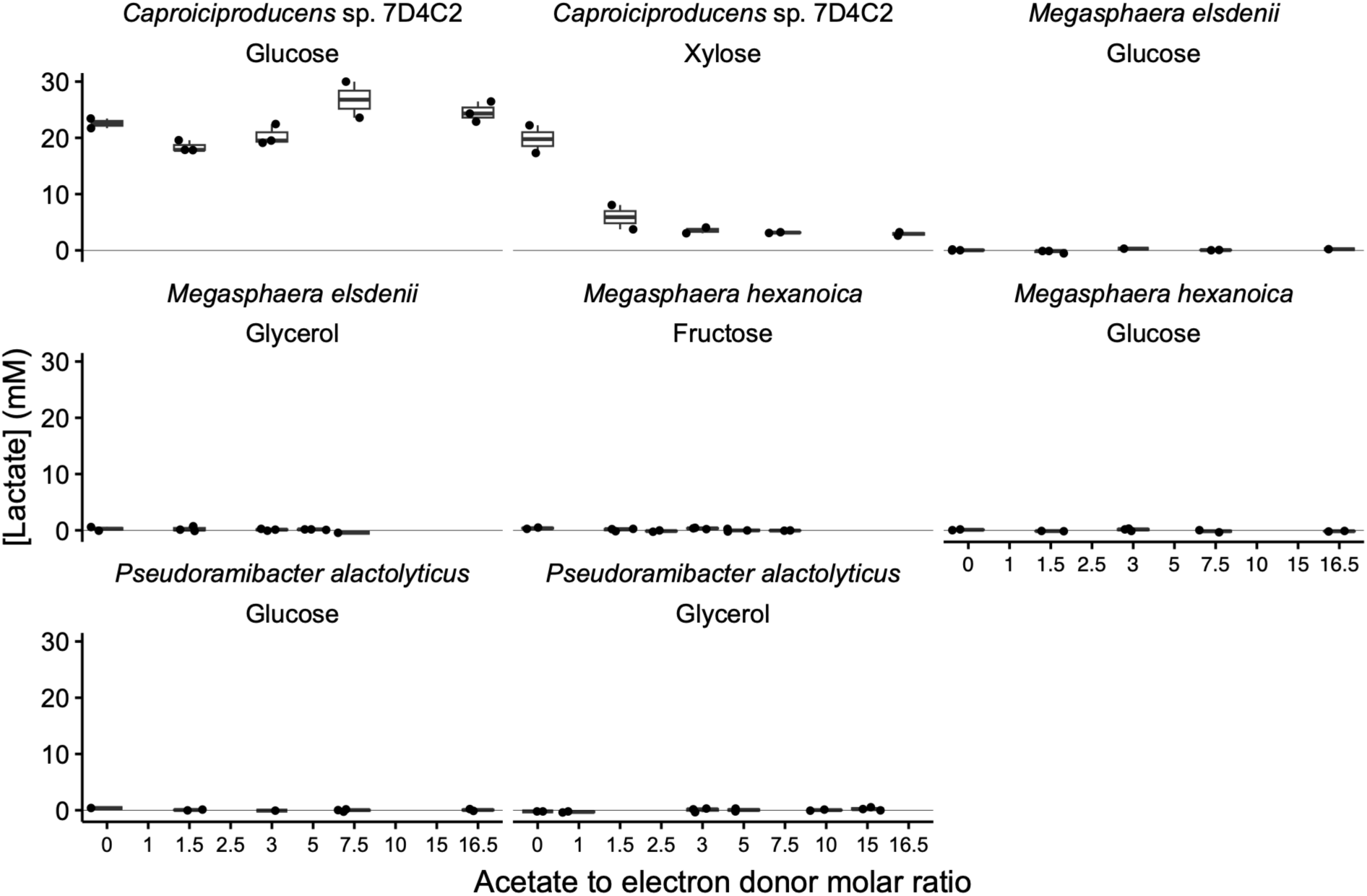
Lactate production of CEB across electron donors and acetate to electron donor ratios.

## References

1. Hackmann, T. J. The vast landscape of carbohydrate fermentation in prokaryotes. FEMS Microbiol Rev 48, fuae016 (2024).

2. Angenent, L. T. et al. Chain Elongation with Reactor Microbiomes: Open-Culture Biotechnology To Produce Biochemicals. Environ. Sci. Technol. 50, 2796–2810 (2016).

3. Prasanna Kumar, D. J., Mishra, R. K., Chinnam, S., Binnal, P. & Dwivedi, N. A comprehensive study on anaerobic digestion of organic solid waste: A review on configurations, operating parameters, techno-economic analysis and current trends. Biotechnol Notes 5, 33–49 (2024).

4. Podolsky, I. A. et al. Harnessing Nature’s Anaerobes for Biotechnology and Bioprocessing. Annu. Rev. Chem. Biomol. Eng. 10, 105–128 (2019).

5. Garcia-Ochoa, F. & Gomez, E. Bioreactor scale-up and oxygen transfer rate in microbial processes: An overview. Biotechnology Advances 27, 153–176 (2009).

6. Weusthuis, R. A., Lamot, I., van der Oost, J. & Sanders, J. P. M. Microbial production of bulk chemicals: development of anaerobic processes. Trends in Biotechnology 29, 153–158 (2011).

7. Rinke, C. et al. Insights into the phylogeny and coding potential of microbial dark matter. Nature 499, 431–437 (2013).

8. Rappé, M. S. & Giovannoni, S. J. The Uncultured Microbial Majority. Annual Review of Microbiology 57, 369–394 (2003).

9. Tramontano, M. et al. Nutritional preferences of human gut bacteria reveal their metabolic idiosyncrasies. Nat Microbiol 3, 514–522 (2018).

10. Calero, P. & Nikel, P. I. Chasing bacterial chassis for metabolic engineering: a perspective review from classical to non-traditional microorganisms. Microb Biotechnol 12, 98–124 (2018).

11. Scarborough, M. J., Lawson, C. E., DeCola, A. C. & Gois, I. M. Microbiomes for sustainable biomanufacturing. Current Opinion in Microbiology 65, 8–14 (2022).

12. Dyussekenova, D. et al. Long-Term Production and Recovery of Medium-Chain Carboxylates from Source-Separated Organics. ACS EST Eng. 6, 2592–2605 (2026).

13. Hackmann, T. J. & Zhang, B. The phenotype and genotype of fermentative prokaryotes. Science Advances 9, eadg8687 (2023).

14. Barker, H. A. & Taha, S. M. Clostridium kluyverii, an Organism Concerned in the Formation of Caproic Acid from Ethyl Alcohol. J Bacteriol 43, 347–363 (1942).

15. Agler, M. T., Wrenn, B. A., Zinder, S. H. & Angenent, L. T. Waste to bioproduct conversion with undefined mixed cultures: the carboxylate platform. Trends in Biotechnology 29, 70–78 (2011).

16. Agler, M. T., Spirito, C. M., Usack, J. G., Werner, J. J. & Angenent, L. T. Chain elongation with reactor microbiomes: upgrading dilute ethanol to medium-chain carboxylates. Energy Environ. Sci. 5, 8189–8192 (2012).

17. Jeon, B. S., Kim, S. & Sang, B.-I. Megasphaera hexanoica sp. nov., a medium-chain carboxylic acid-producing bacterium isolated from a cow rumen. International Journal of Systematic and Evolutionary Microbiology 67, 2114–2120 (2017).

18. Kim, B.-C. et al. Caproiciproducens galactitolivorans gen. nov., sp. nov., a bacterium capable of producing caproic acid from galactitol, isolated from a wastewater treatment plant. International Journal of Systematic and Evolutionary Microbiology 65, 4902–4908 (2015).

19. Lindner, B. G., Nallasamy, D. K., Roberts, L., Gois, I. M. & Lawson, C. E. Coupling Stickland fermentation with reverse β-oxidation represents a new mode of medium-chain carboxylate production. 2026.09.09.750506 Preprint at 10.64898/2026.09.09.750506 (2026).

20. Enabling high-throughput biology with flexible open-source automation. Molecular Systems Biology 17, e9942 (2021).

21. Tanner, R. S., Miller, L. M. & Yang, D. Clostridium ljungdahlii sp. nov., an Acetogenic Species in Clostridial rRNA Homology Group I. International Journal of Systematic and Evolutionary Microbiology 43, 232–236 (1993).

22. Müller, P. et al. High-throughput anaerobic screening for identifying compounds acting against gut bacteria in monocultures or communities. Nat Protoc 1–32 (2023) doi:10.1038/s41596-023-00926-4.

23. King, R. D. et al. The Automation of Science. Science 324, 85–89 (2009).

24. Wang, H., Zhou, W., Gao, J., Ren, C. & Xu, Y. Revealing the Characteristics of Glucose- and Lactate-Based Chain Elongation for Caproate Production by Caproicibacterium lactatifermentans through Transcriptomic, Bioenergetic, and Regulatory Analyses. mSystems e00534–22 (2022) doi:10.1128/msystems.00534-22.

25. Esquivel-Elizondo, S. et al. The Isolate Caproiciproducens sp. 7D4C2 Produces n-Caproate at Mildly Acidic Conditions From Hexoses: Genome and rBOX Comparison With Related Strains and Chain-Elongating Bacteria. Frontiers in Microbiology 11, (2021).

26. Bornstein, B. T. & Barker, H. A. The Energy Metabolism of Clostridium kluyveri and the synthesis of fatty acids. Journal of Biological Chemistry 172, 659–669 (1948).

27. Gois, I. M., Bowers, C. M., Kim, B.-C., Flick, R. & Lawson, C. E. Acetate utilization strategy in chain-elongating bacteria determines butyrate versus medium-chain carboxylate production. Nat Microbiol 11, 1639–1651 (2026).

28. Kang, S., Kim, H., Seung Jeon, B., Choi, O. & Sang, B.-I. Chain elongation process for caproate production using lactate as electron donor in Megasphaera hexanoica. Bioresource Technology 126660 (2021) doi:10.1016/j.biortech.2021.126660.

29. Forsberg, C. W. Nutritional characteristics of Megasphaera elsdenii. Can. J. Microbiol. 24, 981–985 (1978).

30. Dellomonaco, C., Clomburg, J. M., Miller, E. N. & Gonzalez, R. Engineered reversal of the β-oxidation cycle for the synthesis of fuels and chemicals. Nature 476, 355–359 (2011).

31. Duncan, S. H., Hold, G. L., Barcenilla, A., Stewart, C. S. & Flint, H. J. Roseburia intestinalis sp. nov., a novel saccharolytic, butyrate-producing bacterium from human faeces. International Journal of Systematic and Evolutionary Microbiology 52, 1615–1620 (2002).

32. Hillman, E. T. et al. Comparative genomics of the genus Roseburia reveals divergent biosynthetic pathways that may influence colonic competition among species. Microb Genom 6, mgen000399 (2020).

33. Flaiz, M. et al. Caproicibacter fermentans gen. nov., sp. nov., a new caproate-producing bacterium and emended description of the genus Caproiciproducens. International Journal of Systematic and Evolutionary Microbiology 70, 4269–4279 (2020).

34. Gu, Y. et al. Reconstruction of xylose utilization pathway and regulons in Firmicutes. BMC Genomics 11, 255 (2010).

35. Royce, L. A., Liu, P., Stebbins, M. J., Hanson, B. C. & Jarboe, L. R. The damaging effects of short chain fatty acids on Escherichia coli membranes. Appl Microbiol Biotechnol 97, 8317–8327 (2013).

36. Balderrama-Gómez, A. et al. Using Caprylic Acid for the Prevention and Treatment of Helicobacter pylori Infection and Gastric Cancer: A Review. Metabolites 15, 629 (2025).

37. Zhu, X. et al. Microbial Ecological Mechanism for Long-Term Production of High Concentrations of n-Caproate via Lactate-Driven Chain Elongation. Applied and Environmental Microbiology 87, e03075–20 (2021).

38. Roghair, M. et al. Effect of n-Caproate Concentration on Chain Elongation and Competing Processes. ACS Sustainable Chem. Eng. 6, 7499–7506 (2018).

39. Contreras-Dávila, C. A., Zuidema, N., Buisman, C. J. N. & Strik, D. P. B. T. B. Reactor microbiome enriches vegetable oil with n-caproate and n-caprylate for potential functionalized feed additive production via extractive lactate-based chain elongation. Biotechnol Biofuels 14, 232 (2021).

40. Candry, P. et al. A novel high-throughput method for kinetic characterisation of anaerobic bioproduction strains, applied to Clostridium kluyveri. Sci Rep 8, 9724 (2018).

41. Zhang, H. et al. Acoustic Ejection Mass Spectrometry for High-Throughput Analysis. Anal. Chem. 93, 10850–10861 (2021).

42. Liu, Y., Zhong, B. & Lawal, A. Recovery and utilization of crude glycerol, a biodiesel byproduct. RSC Adv. 12, 27997–28008 (2022).

43. Lipids from food waste as feedstock for biodiesel production: Case Hong Kong. Lipid Technology 26, 206–209 (2014).

44. Duber, A., Zagrodnik, R., Juzwa, W., Gutowska, N. & Oleskowicz-Popiel, P. Simultaneous medium chain carboxylic acids and 1,3-propanediol production in a bioaugmented lactate-based chain elongation induced with glycerol. Bioresource Technology 393, 130123 (2024).

45. Leng, L. et al. Shaping microbial consortia in coupling glycerol fermentation and carboxylate chain elongation for Co-production of 1,3-propanediol and caproate: Pathways and mechanisms. Water Research 148, 281–291 (2019).

46. Allaart, M. T., Stouten, G. R., Sousa, D. Z. & Kleerebezem, R. Product Inhibition and pH Affect Stoichiometry and Kinetics of Chain Elongating Microbial Communities in Sequencing Batch Bioreactors. Front Bioeng Biotechnol 9, 693030 (2021).

47. Jarboe, L. R., Royce, L. A. & Liu, P. Understanding biocatalyst inhibition by carboxylic acids. Front Microbiol 4, 272 (2013).

48. Hermans, D. et al. Intestinal mucus protects *Campylobacter jejuni* in the ceca of colonized broiler chickens against the bactericidal effects of medium-chain fatty acids. Poultry Science 89, 1144–1155 (2010).

49. Zimmermann, J., Kaleta, C. & Waschina, S. gapseq: informed prediction of bacterial metabolic pathways and reconstruction of accurate metabolic models. Genome Biol 22, 81 (2021).

